# Steerable Tape-Spring Needle for Autonomous Sharp Turns Through Tissue

**DOI:** 10.1101/2024.09.22.614377

**Authors:** Omar Abdoun, Davin Tjandra, Katie Yin, Pablo Kurzan, Jessica Yin, Mark Yim

## Abstract

Steerable needles offer a minimally invasive method to deliver treatment to hard-to-reach tissue regions. We introduce a new class of *tape-spring* steerable needles capable of sharp turns ranging from 15 to 150 degrees with a turn radius as low as 3mm, which minimizes surrounding tissue damage. In this work, we derive and experimentally validate a geometric model for our steerable needle design. We evaluate both manual and robotic steering of the needle along a Dubins path in 7 kPa and 13 kPa tissue phantoms, simulating our target clinical application in healthy and unhealthy liver tissue. We conduct experiments to measure needle robustness to stiffness transitions between non-homogeneous tissues. We demonstrate progress towards clinical use with needle tip tracking via ultrasound imaging, navigation around anatomical obstacles, and integration with a robotic autonomous steering system.

## I. Introduction

Steerable needles capable of curved paths have been studied in the robotics community for several decades [1], [2], because of their ability to navigate around anatomical obstacles such as bones and sensitive anatomical structures [3], [4], [5]. Clinically, steerable needles could be useful percutaneously for biopsies and treatment of tumors in the right upper lobe of the liver. This region is challenging to access for straight needles due to its proximity to vital structures (lungs) and its location beneath the rib cage. Steerable needles could enable precise navigation around these structures, reducing the risk of damaging critical organs or blood vessels, and enhancing the effectiveness of procedures such as targeted drug delivery, thermal ablation, or biopsy sampling for liver tumors.

Current approaches without steerable needles primarily utilize straight devices, which can be limited by anatomical obstacles and require more skill to navigate. This increases risk of negative outcomes, such as injuring the lungs and leading to complications such as pneumothorax (collapsed lung), and inaccurate targeting of these lesions can lead to a higher rate of complications and recurrence [6], [7], [8]. This limitation necessitates the development and use of steerable needles that can navigate around critical structures, reducing the likelihood of such adverse events and improving the accuracy, safety, and efficacy of percutaneous procedures.

The robotics research community has focused on developing several key improvements to steerable needle approaches for clinical impact. These include four challenges: 1) the ability to make sharp turns, 2) including more human-in-the-loop control, 3) modeling the mechanics of varied tissue interaction, and 4) minimizing tissue damage.

The ability to make sharp turns is key to maximizing accessible workspace while minimizing tissue damage [9], [10], [2]. Including more human-in-the-loop control will both make it easier to use (intuitive) for the surgeon and reduce cost [11], [12]. A large portion of the cost of robotic steerable needles may come from the robotic drive units (motors, sensors, control computation, and regulated power). A device that can be used both manually and robotically provides greater flexibility in clinical use. Quick and simple procedures such as deep ultrasound-guided steered IV placements can be performed manually, while more complex procedures (e.g., liver tumor ablations) can benefit from robotic control. For the tissue interaction challenge, [13], [14], [15] others have examined the planning and modeling of varied tissues and interactions with steerable needles.

Tape spring steerable needles (Fig. 1) can potentially address the four key challenges. This new class of tape-spring needles have the smallest turning radius that the authors have seen published [16]. Compared to straight needles of similar diameter and length, tape-spring needles add no additional damage in a straight path [17]. In this paper, we show that the needle paths are invariant to tissue stiffness and, as a result, are likely to work well with both direct human-in-the-loop and robotic control. This adaptability allows the device to be matched to the appropriate clinical scenario.

**Fig 1.**
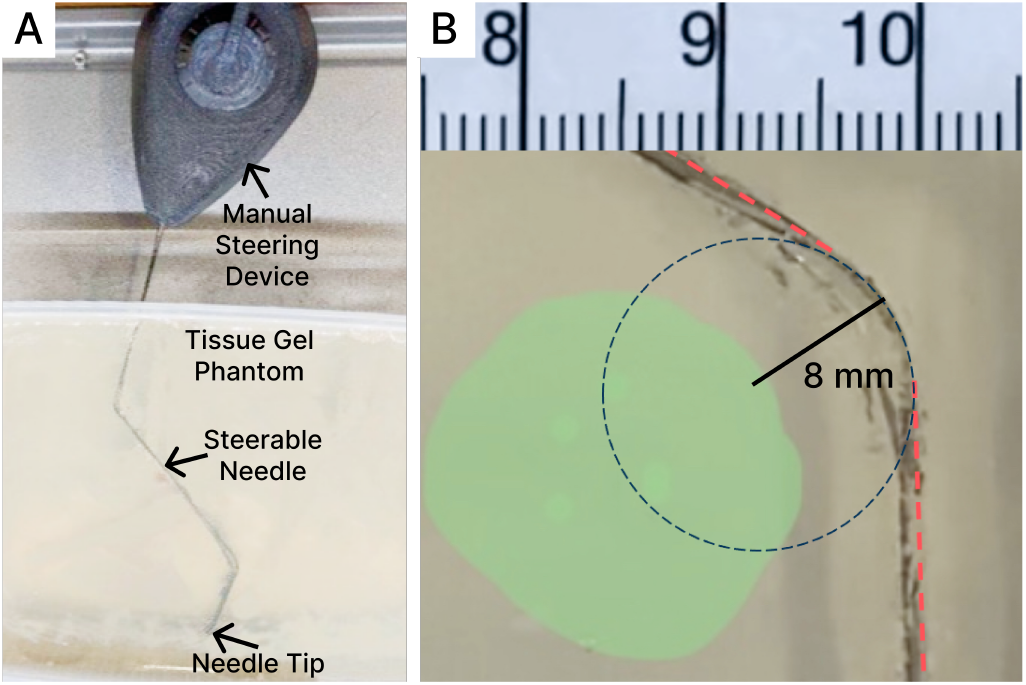
A) We present a tape spring steerable needle, capable of at least two sharp turns in opposing directions, demonstrated in a gel tissue phantom. B) A magnified image of a sharp turn, which forms a 55^*o*^ arc with radius 8 mm, as the needle navigates around an obstacle. Small turn radii are critical for minimizing tissue damage and maximizing the workspace of the needle. We also show how the straight and curved segments of the needle follow Dubins path for intuitive manual and robotic steering.

The main contributions of this paper are as follows:

1) The design and fabrication of a tape-spring steerable needle and robot steering system is presented along with the effectiveness of both **manual and robotic** control measuring the tracking error moving through 7kPa and 13kPa tissue phantoms that simulate our target clinical application in liver tissue.
2) A geometric model of the needle trajectory, derived and experimentally validated, closely follows Dubins path. This indicates compatibility with more complex path-planning algorithms in future work.
3) Autonomous steering, ultrasound needle tracking, and obstacle navigation capabilities of the robotically and manually steered tape-spring needle are demonstrated.

## II. Related Work

The concentric tube steerable needle design [18] (the active cannula class of steerable needle) has been shown in the robotics community as a device that is similar to a robot arm, consisting of a continuum manipulator whose endpoint can be controlled by twisting concentric tubes that have biased bends embedded in them. Bevel tip needles are another class of flexible needle [19] that passively form curved paths as they are inserted. Twisting the needle thus changes the plane of the curve that the needle takes through tissue. Controlling a bevel-tip needle path is not intuitive; for example, to go straight, the needle must be continuously twisted, forming a helix (and increasing tissue damage). Other steerable needle approaches have tubular bodies and vary the elements at the head of the needle. These variations include precurved stylets, programmable bevels, and tips actuated with cables [1]. Each has various performance metrics when evaluating suitability for treatments, including turning radius, tissue damage, and susceptibility to variance in the tissues’ mechanical properties.

Robotic needle steering has the potential to minimize patient tissue damage while making steerable needle procedures more accessible [20]. Due to the manual nature of current protocol, patient outcome and treatment success are subject to the skill and experience of the surgeon. Much of the work in robotic needle steering focuses on variations of bevel-tip needles [21], [22], [23], [24], [25], [26]. Here, the robot steers by controlling insertion and rotational forces at the needle base, typically through linear and rotational motors rigidly affixed to the needle [27]. In some cases, the robot may also control a cable attached to the flexible tip of the needle [28]. Feedback for closed loop control and path planning generally employs computer vision techniques with ultrasound or CT scans to track the needle tip, needle path, target region, and anatomical obstacles [29]. Most robot systems are demonstrated on *ex-vivo* organs or phantom gels that share the mechanical characteristics of the target tissue. Towards convincingly demonstrating the clinical applicability of robotic needle steering [30], [14] notably perform *in-vivo* procedures with autonomous robotic needle steering. We build upon these works by introducing a robot system to autonomously steer a *tape-spring* needle design by controlling insertion velocity and cable displacement of the needle.

## III. Previous Work

In [16], we characterized the bending moment required to induce buckling and presented a model of the forces on the needle being comprised of cutting forces from the tip, friction forces along the sides and stress forces induced from a bend. The latter forces are the ones that impact tissue damage. In addition, we demonstrated ultrasound guided intravenous needle (IV) placement at extreme angles in phantom tissue and an ultrasound guided turn in porcine liver, as well as a novel method for assessing tissue damage [17]. Another particularly useful property for isotropic material tape springs is that when buckled, the buckled portion maintains a constant radius arc of a circle. The radius of this arc has been shown to be nearly the same as the transverse bend radius [31]. This is shown in Fig 2a. In addition, Figure 2b shows a schematic of a tape spring with two turns of equal radius. The “ploy” region is the area where the beam transitions from curved transversely to curved longitudinally and is approximated as being straight [31] with no discontinuities, so that there is no part of the path that is not either straight or the arc of a circle. We have shown that when molding 0.038mm thick 1095 steel tape spring needles with a transverse radius of about 3mm we can achieve turns with radii of 3.3mm in phantom tissue and a device width of only 3mm [16].

**Fig 2.**
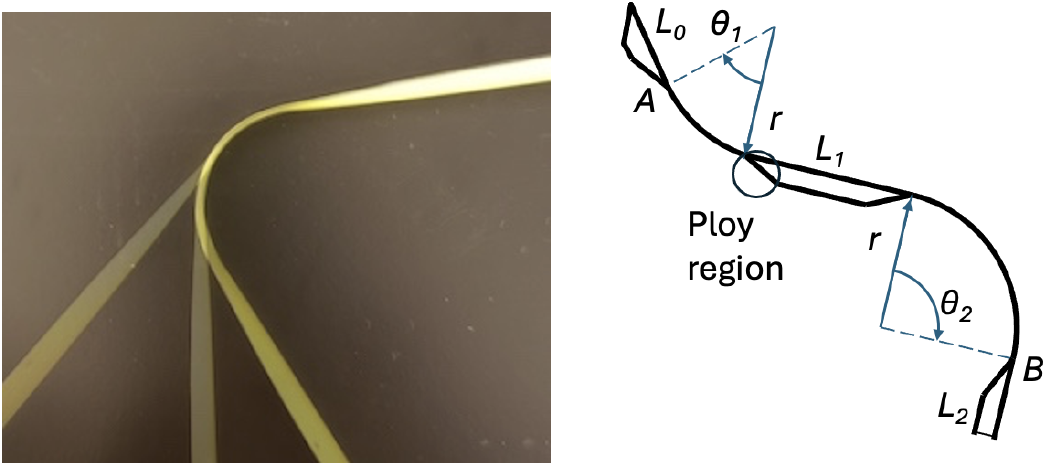
a) Three images of a tape spring in air at three different bend angles forming overlaid showing the same radius arcs of a circle. b) Schematic of a tape spring with two turns. Note that the radii of each turn is the same.

## IV. Design and Fabrication

### A. Steerable Needle

We utilize a tape spring [32], [33], a transversely curved elastic beam commonly used in tape measures, as the underlying mechanism for our steerable needle design. The needle can be steered by inducing buckling near the sharpened tip of the needle, which generates sharp turns about a small turn radius [16]. Tape springs have four key properties that are well suited to steerable needles:

1) When buckled at the head, the beam forms a turn that propagates down the beam as the needle is inserted (which leads to follow-the-leader behavior, forming a clean needle path).
2) The forces required to maintain this buckled state are very low, and if using low damping factor materials, the propagation forces are also low; in total, on the order of 1 mN. This means that the stresses imposed on the surrounding tissue can be small (approximately 7.5 Pa), which reduces tissue damage [17].
3) When straight, the beam can support compressive loads, allowing the needle to puncture tissue and overcome friction forces for deeper paths, even with a small mm-scale cross-sectional profile.
4) Since the needle turning behavior does not depend on a passive geometry of the cutting portion (like bevel-tip) and the forces on tissues for turning are low, the effect of tissue variance on steering ability is low (Section IVA and B).

To fabricate our tape-spring needle, we use a water jet to cut the needle from a sheet of 50 *µ*m bio-compatible 1095 steel and post-process it with a guillotine to smooth the edges, reducing friction during tissue insertion. The needle pattern features a pointed tip, a narrow neck intended to buckle, and holes for cable routing (Fig. 3A, B). A 2-part aluminum mold, consisting of a 6.4 mm diameter cylindrical cutout and solid rod creates the transverse curve of the needle. The needle and mold are heated to 260^°^C for one hour and quenched in room-temperature water. This process is performed three times to reduce spring back and ensure the desired needle curvature. After removing the needle from the mold, cables are routed through the holes and attached at the tip with cyanoacrylate adhesive.

**Fig 3.**
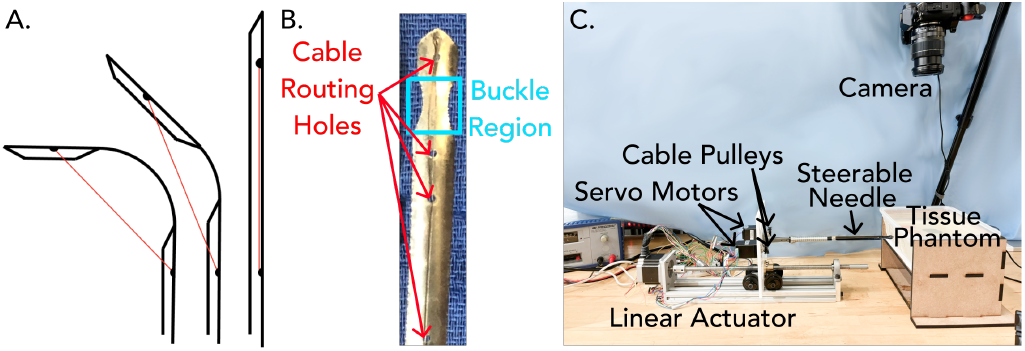
A. Cable-induced buckling near the needle tip enables steering. The intended buckling region is located between the cable routing holes. B. Photo of fabricated tape-spring needle, showing needle tip geometry, buckle region, and cable routing holes. C. System diagram of autonomous steering robot.

### B. Robot System for Autonomous Steering

We design our robot steering system to achieve the following goals: 1) control needle insertion velocity, 2) precisely modulate cable displacement for steering, and 3) track the needle tip with camera feedback (Fig. 3C). We control the needle insertion velocity using a linear actuator with a NEMA 23 stepper motor (200 steps) and a T8×2 lead screw (0.01 mm resolution). The lead screw drives a laser-cut acrylic plate and aluminum slider carriage. The needle, 3D-printed spool, cable guide ring, and two NEMA 11 stepper motors are mounted to the acrylic plate. The NEMA 11 stepper motors provide 9.5 N·cm of torque and 200 steps per revolution (1.8 degrees per step) to wind the cable for steering. The attached pulleys have a 15.6 mm diameter, translating to 0.245 mm of cable displacement per step.

For insertion into tissue phantoms, the needle moves forward with a constant velocity of 0.5 mm/sec. We steer the needle tip by controlling cable displacement with the pulleys. The combination of cable displacement and linear movement allows us to set the turn angle. A spring sheath holds the needle’s base to prevent buckling in unintended areas. A camera provides feedback to enable closed-loop control and obstacle detection.

## V. System Modeling and Characterization

### A. Needle Turn Model and Validation

#### 1) Geometric Model for Controlled Needle Turns

Intuitive control of the tape-spring needle (by pulling on the cables) requires an intuitive mapping from the cable displacement (CD) to the needle tip turn angle. For the mapping to be intuitive, the relationship between control input (cable displacement) and output (needle tip turn angle) should be monotonic, smooth, and ideally linear. An example of *intuitive mapping* is a car’s steering wheel relationship to the car’s turning angle. Most modern cars use Ackermann steering [34] which has a roughly linear relationship between steering wheel angle and the car’s turning angle near 0 (going straight) but becomes very non-linear at extreme angles.

Below, we derive a geometric model that relates cable displacement to needle turn angle. We use this model to guide the control parameters of motion primitives used for our low-level autonomous steering controller.

The theoretical mapping of CD to the steering angle can be derived from analyzing the geometry at the head of the needle. Figure 3a shows the head of the needle where a cable attaches from the bottom of the head to the tip of the head. Pulling on this cable pulls the tip down. Bends initiate at the midpoint between the cable entry holes. This arc of fixed radius grows from this point out as the bend increases. The length of tape spring between the cable entry holes is *TSL* (Total Segment Length).

*TSL* can also geometrically be described as the sum of the lengths of the straight segments between the holes, distal length, *DL*, proximal length, *PL*, and the arc length, *AL*, of the bent segment as can be seen in Figure 4.

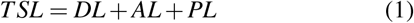

**Fig 4.**
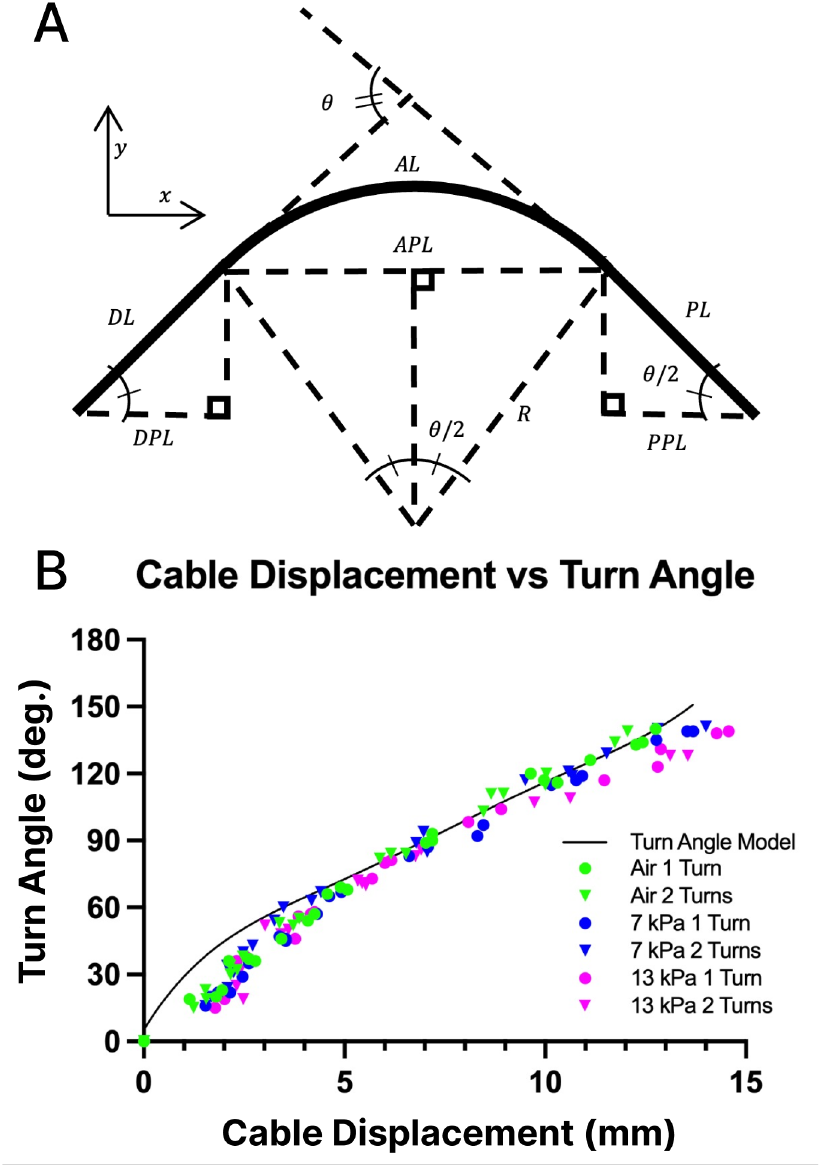
A) Schematic of the proposed geometric turn angle model with labeled geometry between the cable entry holes of length *TSL*. Note the bend is symmetric. B) Experimental validation of our geometric model. We map cable displacement to needle turn angle for both single and double turns in air, 7 kPa gel, and 13 kPa gel.

*CD* can be represented as the difference between the *TSL* and the cable length, *CL*, that runs straight between the two holes, as in Equation 2.

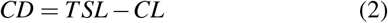

Equation 3 represents *CL* as the projections of *DL, PL*, and *AL* in the *x* direction, which are *DPL, PPL*, and *APL*, respectively.

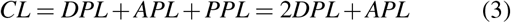

By design symmetry, *DL* and *PL* are equal in length and are solved for in Equation 4 by substitution and rearranging Equation 1.

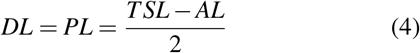

*AL* is derived by geometry as the arc length is the product of the angle and radius *R. DPL, PPL*, and *APL* are solved for in Equations 5 and 6 using the geometry of right triangles.

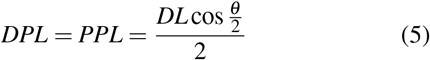

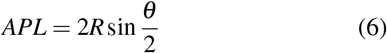

Substituting Equations 5 and 6 into Equation 3 yields Equation 7.

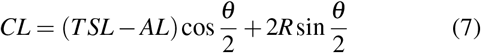

Substituting Equation 7 into Equation 2 and rearranging yield the relationship between *CD* and *θ* in Equation 8

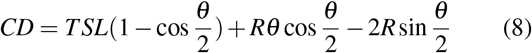

In order to determine the amount of cable pull required to turn a given amount Equation 8 can be inverted using standard numerical techniques such as Newton-Raphson.

#### 2) Experimental Validation

The prototype needle was inserted into 7 kPa and 13 kPa stiffness materials and in air with increasing bend angle. We use video processing to extract the length of cable pulled to induce the bend and the resulting turn angle. A second bend was introduced in the reverse direction with the angles and cable pull recorded.

Figure 4 shows the angle vs cable displacement for three different stiffness conditions and two different numbers of turns. In all cases, the data matches fairly closely with the theoretical model expressed in Equation 8.

The theoretical model is purely kinematic and does not include any cable stretch, slack, or deflections in the body of the needle. At small cable displacements the initial moment required to induce a buckle results in motion of the cable without concomitant changes in the angle, which can be seen in the figure at CD of less than 3mm. This slack in the cable can be minimized by pre-tensioning the cables or with force control in the future. The effect of slack could reasonably explain the difference between the model and experimental data seen at less than 3mm.

The match of all data points with Equation 8 is a strong indicator that the tape spring needle is immune to stiffness changes from 7kPa to 13kPa. At larger stiffness, for example in cirrhotic livers, it is not as clear that this immunity will remain and would be worthwhile for future exploration.

### B. Quantifying Needle Trajectory Error

Low tracking error in the needle trajectory is critical to prevent unintended damage to vital structures. Here, we quantify the tracking error of both robotic and manual needle steering along a pre-planned Dubins path through 7 kPa (healthy liver) and 13 kPa (unhealthy liver) tissue phantom gels. We pattern a transparent stencil with a Dubins path consisting of a 5 cm straight segment, a 30° turn with an 8 mm radius, and a 3 cm post-turn straight segment. The stencil is overlaid on the gel. We *manually and robotically* steer the needle approximately 1.5 cm below the surface, following the stencil path as closely as possible.

We analyzed key frames in videos of the needle’s trajectory with ImageJ software. We define two performance metrics: 1) The deviation distance of the needle tip from the stencil at each centimeter marking, and 2) the deviation area of the needle’s final path from the stencil path. The area of deviation was divided to calculate the offset of the needle from the stencil in millimeters (Table I). In addition, the error value at select points along the path in both substrate stiffnesses with both manual and robotic control can be seen in Fig. 5.

**Fig 5.**
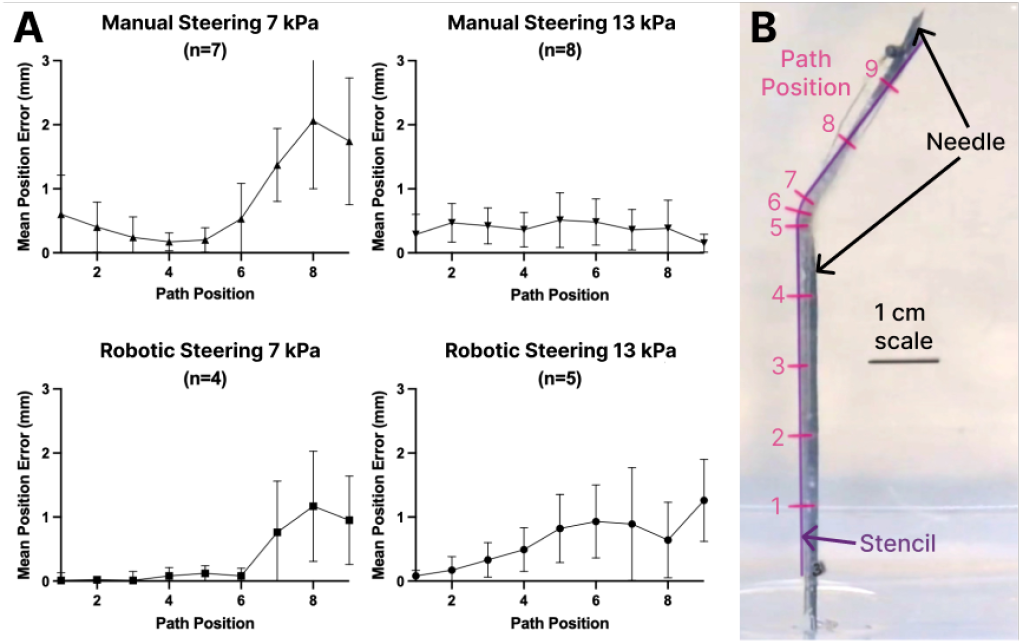
A) Mean path error at select points along the target trajectory with both manual and robotic insertions across 7kPa and 13kPa gels, showing low placement error across all trials. B) Steerable needle closely following target Dubins path with measurement points labeled in red.

**TABLE I.** Mean needle path error from intended Dubins path. The refraction error at 1.5 cm depth in the gel is 0.1 mm and the pixel density of our image data is 11.5 pixels per 1mm.

| Trial | Path Error (mm) | Std. Deviation (mm) |
| --- | --- | --- |
| 7 kPa Manual (n=8) | 0.48 | 0.41 |
| 7 kPa Robotic (n=4) | 0.12 | 0.06 |
| 13 kPa Manual (n=8) | 0.21 | 0.17 |
| 13 kPa Robotic (n=5) | 0.16 | 0.10 |

Navigating through the 13 kPa gel demonstrated less variability and greater accuracy in following the planned path than the 7 kPa gel. However, the average path error was low for both the manual and the robotic insertions. In some of the trials, there is a larger error near the end of the path than there is at the beginning. This may be due to accumulated error and future work can include improvement of the steering algorithm to address this. Overall, the success in following Dubins path with low deviation in both substrate stiffnesses demonstrates that the needle can be controlled using the same steering algorithm and inputs across a range of clinically relevant tissue mechanical properties. Critically, all modalities and substrate stiffnesses had mean position error less than the 2.7 mm allowable needle placement error of surveyed Interventional Radiologists [35]. In addition, comparable accuracy of both manual and robotic control shows that the needle can be used with either steering method without trade-off in accuracy.

### C. Incidence Angle Robustness

A concern from needle interaction with tissue occurs at the interface of different materials (for example going from healthy to diseased liver) where the path of the needle may diverge from the ideal modeled path. If the angle of incidence while entering the higher-stiffness material occurs at a glancing angle, there can be large errors in needle path. Errors of 1 cm have been reported in [2] when interacting with a “tissue membrane” that illustrates this case.

Here, we evaluate robustness of the steerable needle in executing a planned path *across* substrates of different stiffnesses at an acute incidence angle (Fig. 6). We fill containers with two distinct sections of gel: one section of 7 kPa gel (clear) and another of 13 kPa gel (pink). The sections are pressed together to create 10°, 20°, or 30° incidence angles. We manually steer the needle in the 7 kPa gel and 13 kPa gel following a straight trajectory, with nine total insertions for each transition angle. We analyze an image of the needle path, measuring the angle of the needle using ImageJ’s angle tool. The pixel density of these images are 60 pixels/cm and we quantified the precision of these manual measurements as 0.03°. The angle measurements are calculated with the entry point into the gel, the point of transition between the two gels, and the tip of the needle. The target angle across the transition was 180° and the results of this experiment can be seen in Table II.

**Fig 6.**
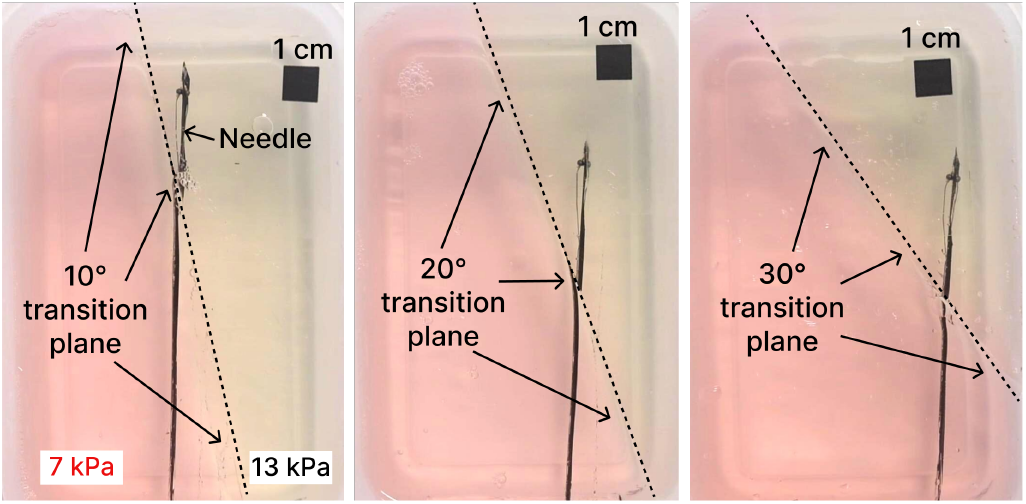
Steerable needles navigating from low stiffness (7 kPa)to high stiffness (13 kPa) gels at 10°, 20°, and 30° of incidence with minimal effect on target path.

**TABLE II.**
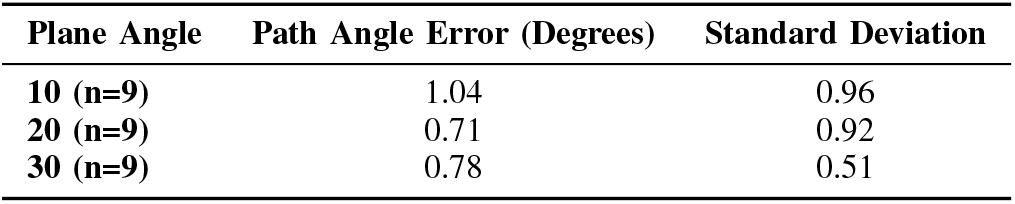
Mean needle angle error across incidence angles between 7 kPa (healthy liver stiffness) and 13 kPa (unhealthy liver stiffness) gels. The pixel density of our image data is 6 pixels/mm and the angle precision is 0.03^°^.

| Plane Angle | Path Angle Error (Degrees) | Standard Deviation |
| --- | --- | --- |
| 10 (n=9) | 1.04 | 0.96 |
| 20 (n=9) | 0.71 | 0.92 |
| 30 (n=9) | 0.78 | 0.51 |

All incidence angles tested showed minimal effect on the intended needle trajectory. These low degrees of error demonstrate that the needle was able to follow the planned straight trajectory when transitioning between gels of different stiffnesses. This is another indication that the needle steering will not be overly affected by tissue properties. In the future, the full range of compatible tissue mechanical properties can be characterized.

## VI. Demonstrations

### A. Anatomical Obstacle Avoidance

This demonstration simulates a clinical procedure using a steerable needle to access a tumor in the upper right lobe of the liver (Fig. 7). This region is challenging to reach because the needle must navigate around the ribs and lungs, risking lung puncture or significant tissue damage.

**Fig 7.**
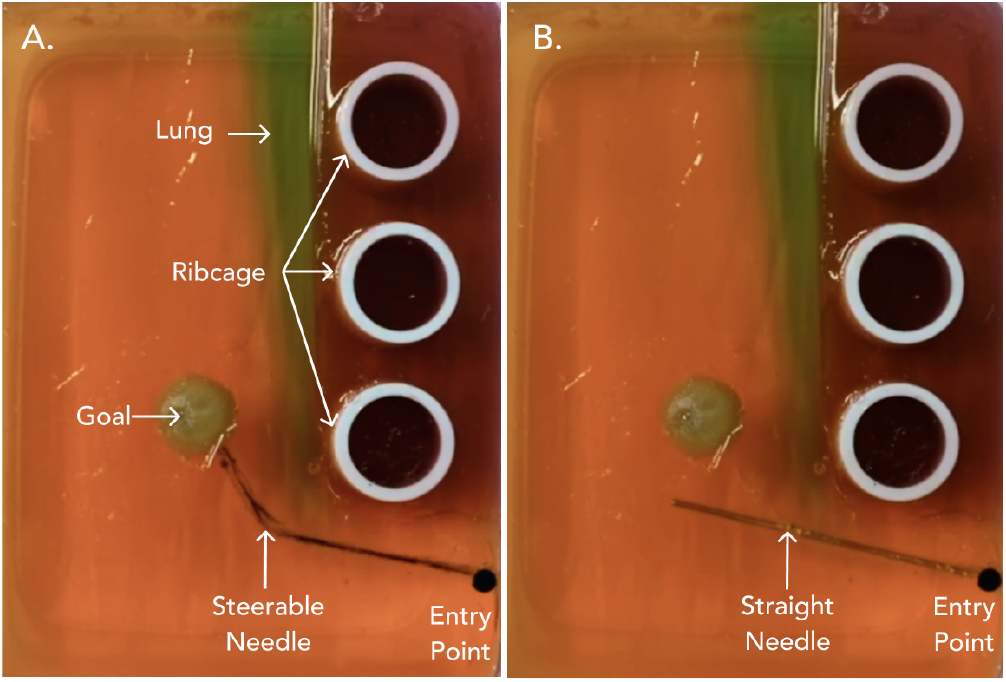
A. We demonstrate the tape-spring steerable needle navigating around anatomical obstacles, such as the lungs and ribs, to reach the goal region in the liver. B. In contrast, a straight needle is unable to reach the goal.

We show that we can manually steer the tape-spring needle to navigate from the entry point, around the lungs and rib bones to the tumor in the liver. In contrast, a straight needle is unable to reach the same goal region.

### B. Ultrasound Needle Tracking

Ultrasound is a widely used imaging modality that provides real-time visualization of needle insertions into tissue. Many procedures involving needle interventions are ultrasound-guided, such as biopsies, drug delivery, or catheter insertions. This demonstration shows that the steerable needle is compatible with ultrasound imaging. Furthermore, the ultrasound image can be processed to track the needle tip for closed loop control.

In the ultrasound image, the needle is hyperechoic and appears as bright white while the surrounding phantom tissue is hypoechoic (black). The extra white reflection artifacts visible in the ultrasound image are due to reflections in the phantom tissue and can be thresholded out of the image for accurate tip tracking.

### C. Autonomous Robotic Needle Steering

The autonomous robot steering controller consists of: 1) a high level controller for obstacle detection and needle tip tracking feedback, and 2) a low level controller to command the motors for insertion velocity and cable displacement. The high level controller is implemented in Python and uses color thresholding (OpenCV) to identify anatomical obstacles, the goal region, and the needle tip. The obstacles and goal are identified prior to inserting the needle and assumed to be constant throughout the needle trajectory. The high level controller then generates a collision-free needle trajectory using Dubins path, assuming that the needle has two motion primitives: a straight step (1 cm) and a 30 degree right turn. Using serial communication, the high level controller commands the low level controller (implemented with Arduino C++) to execute the desired motion primitive. After the motion primitive is executed, the needle tip position is compared against the desired needle path. If the actual needle position deviates from the path such that a collision will occur or the needle can no longer reach the goal, the high level controller withdraws the needle for another attempt.

We demonstrate our autonomous steering system with a theoretical clinical scenario: navigating between two blood vessels such as the portal circulation and hepatic circulation, to reach a liver tumor (Fig. 9). The obstacles and goal region are identified prior to needle insertion. The robot steering system generates the path and successfully executes the planned trajectory to reach the goal.

**Fig 8.**
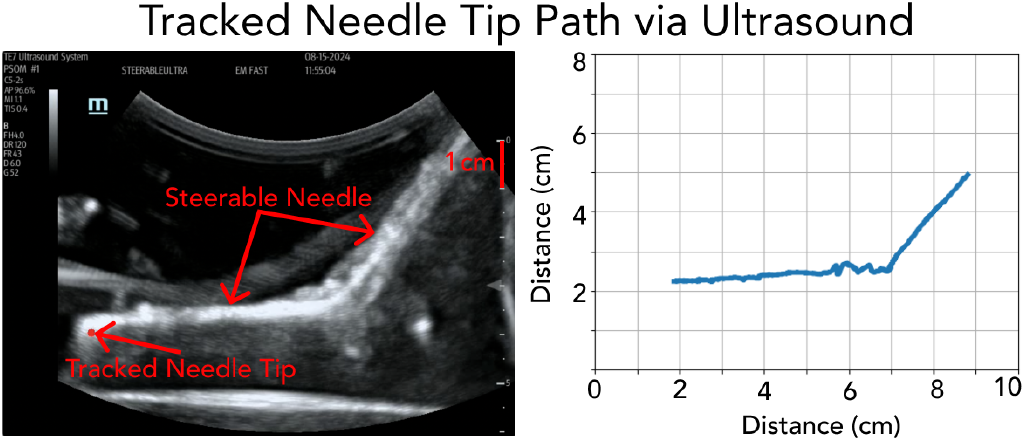
The tape-spring needle tip can be tracked via ultrasound for closed-loop feedback, the most common imaging modality for relevant clinical procedures. From the video taken on the ultrasound machine, needle tip tracking using OpenCV was employed and the path of the needle tip trajectory was plotted [right] using Matplotlib.

**Fig 9.**
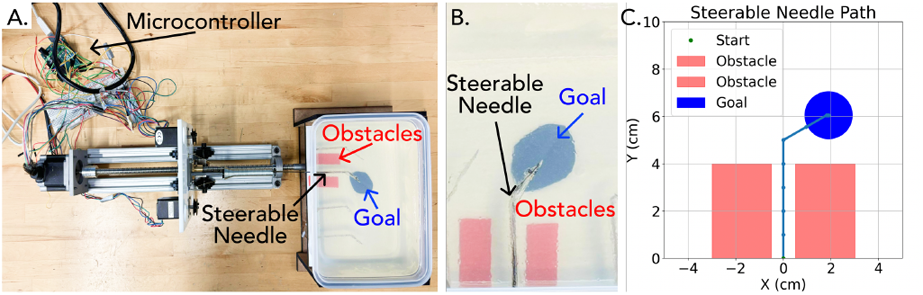
Autonomous needle steering demo. A) Robot system used to steer needle. B) Photo of anatomical obstacles (blood vessels), steerable needle, and goal region. C) Collision-free Dubins path for steerable needle generated by autonomous steering system.

## VII. Conclusions AND Future Work

In this paper, we have demonstrated the capabilities of the tape spring steerable needle device, highlighting its small turn radius and the ability to be accurately controlled both robotically and manually. Our experiments show that the device achieves path position errors averaging less than 0.5 mm. We developed a geometric model to use as an input for steering, which was experimentally validated to demonstrate that the needle’s trajectory closely follows Dubins paths, facilitating intuitive robotic steering. The autonomous steering demonstrations highlight the capabilities of the device for closed-loop control. We have also demonstrated the device’s compatibility with ultrasound and its ability to track the needle tip under ultrasound guidance using computer vision. These results indicate that this device could be applied to a variety of clinical applications due to its simple and predictable control that allows for both manual and robotic steering. This work builds on prior research demonstrating the device’s minimal tissue damage to surrounding tissues and its development of a dynamic force model. It also retains the smallest turn radius of any steerable device documented. Future work will focus on expanding the steering capabilities of the needle into 3D, leveraging the torsional properties of tape springs to enhance maneuverability. Further development will include improving the turning algorithm to minimize error, particularly at the end of the path. Additionally, we aim to characterize the device’s performance at more varied angles.

## Supporting information

Demo Video Included with Manuscript Submission

## Acknowledgements

We thank Jeremy Wang, Britny Major, and Megan Santamore for fabrication and design assistance and Eza Koch and Elizabeth Tu for their work in early prototyping of this device.

## References

[1] N. J. Van De Berg, D. J. Van Gerwen, J. Dankelman, and J. J. Van Den Dobbelsteen, “Design choices in needle steering—a review,” IEEE/ASME Transactions on Mechatronics, vol. 20, no. 5, pp. 2172– 2183, 2014.

[2] K. B. Reed, A. Majewicz, V. Kallem, R. Alterovitz, K. Goldberg, N. J. Cowan, and A. M. Okamura, “Robot-assisted needle steering,” IEEE robotics & automation magazine, vol. 18, no. 4, pp. 35–46, 2011.

[3] Y. Duan, J. Ling, Z. Feng, T. Ye, T. Sun, and Y. Zhu, “A survey of needle steering approaches in minimally invasive surgery,” Annals of Biomedical Engineering, vol. 52, no. 6, p. 1492–1517, Mar 2024.

[4] M. Scali, T. P. Pusch, P. Breedveld, and D. Dodou, “Needle-like instruments for steering through solid organs: A review of the scientific and patent literature,” Proceedings of the Institution of Mechanical Engineers, Part H: Journal of Engineering in Medicine, vol. 231, no. 3, pp. 250–265, 2017, pMID: 28056627. [Online]. Available: 10.1177/0954411916672149

[5] Y. Z. Kaiyu Wu, Bing Li and X. Dai, “Review of research on path planning and control methods of flexible steerable needle puncture robot,” Computer Assisted Surgery, vol. 27, no. 1, pp. 91–112, 2022, pMID: 36052822. [Online]. Available: 10.1080/24699322.2021.2023647

[6] A. Kambadakone, V. Baliyan, H. Kordbacheh, R. N. Uppot, A. Thabet, D. A. Gervais, and R. S. Arellano, “Imaging guided percutaneous interventions in hepatic dome lesions: Tips and tricks,” World Journal of Hepatology, vol. 9, no. 19, pp. 840–849, Jul 2017.

[7] S. A. Curley, P. Marra, K. Beaty, L. M. Ellis, J.-N. Vauthey, E. K. Abdalla, and et al., “Early and late complications after radiofrequency ablation of malignant liver tumors in 608 patients,” Annals of Surgery, vol. 239, no. 4, pp. 450–458, 2004.

[8] S. Mulier, Y. Ni, J. Jamart, T. Ruers, G. Marchal, and L. Michel, “Local recurrence after hepatic radiofrequency coagulation: multivariate meta-analysis and review of contributing factors,” Annals of Surgery, vol. 242, no. 2, pp. 158–171, 2005.

[9] P. J. Swaney, J. Burgner, H. B. Gilbert, and R. J. Webster, “A flexure-based steerable needle: high curvature with reduced tissue damage,” IEEE Transactions on Biomedical Engineering, vol. 60, no. 4, pp. 906–909, 2012.

[10] T. K. Adebar, J. D. Greer, P. F. Laeseke, G. L. Hwang, and A. M. Okamura, “Methods for improving the curvature of steerable needles in biological tissue,” IEEE Transactions on Biomedical Engineering, vol. 63, no. 6, pp. 1167–1177, 2015.

[11] Z. Wang, I. Reed, and A. M. Fey, “Toward intuitive teleoperation in surgery: Human-centric evaluation of teleoperation algorithms for robotic needle steering,” in 2018 IEEE International Conference on Robotics and Automation (ICRA). IEEE, 2018, pp. 5799–5806.

[12] J. M. Romano, R. J. Webster, and A. M. Okamura, “Teleoperation of steerable needles,” in Proceedings 2007 IEEE International Conference on Robotics and Automation. IEEE, 2007, pp. 934–939.

[13] R. Alterovitz, A. Lim, K. Goldberg, G. S. Chirikjian, and A. M. Okamura, “Steering flexible needles under markov motion uncertainty,” in 2005 IEEE/RSJ International Conference on Intelligent Robots and Systems. IEEE, 2005, pp. 1570–1575.

[14] A. Majewicz, S. P. Marra, M. G. Van Vledder, M. Lin, M. A. Choti, D. Y. Song, and A. M. Okamura, “Behavior of tip-steerable needles in ex vivo and in vivo tissue,” IEEE Transactions on Biomedical Engineering, vol. 59, no. 10, pp. 2705–2715, 2012.

[15] S. Misra, K. Ramesh, and A. M. Okamura, “Modeling of tool-tissue interactions for computer-based surgical simulation: A literature review,” Presence, vol. 17, no. 5, pp. 463–491, 2008.

[16] O. T. Abdoun and M. Yim, “A tape spring steerable needle capable of sharp turns,” bioRxiv, doi:10.1101/2023.05.04.539394, [online] https://www.biorxiv.org/content/early/2023/05/07/2023.05.04.539394, 2023. [Online]. Available: https://www.biorxiv.org/content/early/2023/05/07/2023.05.04.539394

[17] O. T. Abdoun and M. Yim, “Assessing tissue damage around a tape spring steerable needle with sharp turn radii,” IEEE Open Journal of Engineering in Medicine and Biology, 2024.

[18] H. B. Gilbert, D. C. Rucker, and R. J. Webster III, “Concentric tube robots: The state of the art and future directions,” in Robotics Research: The 16th International Symposium ISRR. Springer, 2016, pp. 253–269.

[19] R. J. Webster III, J. S. Kim, N. J. Cowan, G. S. Chirikjian, and A. M. Okamura, “Nonholonomic modeling of needle steering,” The International Journal of Robotics Research, vol. 25, no. 5-6, pp. 509– 525, 2006.

[20] M. Babaiasl, F. Yang, and J. P. Swensen, “Robotic needle steering: state-of-the-art and research challenges,” Intelligent Service Robotics, vol. 15, no. 5, pp. 679–711, 2022.

[21] R. Alterovitz, K. Goldberg, and A. Okamura, “Planning for steerable bevel-tip needle insertion through 2d soft tissue with obstacles,” in Proceedings of the 2005 IEEE international conference on robotics and automation. IEEE, 2005, pp. 1640–1645.

[22] R. Seifabadi, E. E. Gomez, F. Aalamifar, G. Fichtinger, and I. Iordachita, “Real-time tracking of a bevel-tip needle with varying insertion depth: Toward teleoperated mri-guided needle steering,” in 2013 IEEE/RSJ International Conference on Intelligent Robots and Systems. IEEE, 2013, pp. 469–476.

[23] A. Favaro, A. Segato, F. Muretti, and E. De Momi, “An evolutionary-optimized surgical path planner for a programmable bevel-tip needle,” IEEE Transactions on Robotics, vol. 37, no. 4, pp. 1039–1050, 2021.

[24] A. Yamada, S. Naka, N. Nitta, S. Morikawa, and T. Tani, “A loop-shaped flexible mechanism for robotic needle steering,” IEEE Robotics and Automation Letters, vol. 3, no. 2, pp. 648–655, 2017.

[25] S. Misra, K. B. Reed, A. S. Douglas, K. Ramesh, and A. M. Okamura, “Needle-tissue interaction forces for bevel-tip steerable needles,” in 2008 2nd IEEE RAS & EMBS international conference on biomedical robotics and biomechatronics. IEEE, 2008, pp. 224–231.

[26] R. J. Webster, J. Memisevic, and A. M. Okamura, “Design considerations for robotic needle steering,” in Proceedings of the 2005 IEEE International Conference on Robotics and Automation. IEEE, 2005, pp. 3588–3594.

[27] G. Gerboni, J. D. Greer, P. F. Laeseke, G. L. Hwang, and A. M. Okamura, “Highly articulated robotic needle achieves distributed ablation of liver tissue,” IEEE robotics and automation letters, vol. 2, no. 3, pp. 1367–1374, 2017.

[28] B. Konh, B. Padasdao, Z. Batsaikhan, and J. Lederer, “Steering a tendon-driven needle in high-dose-rate prostate brachytherapy for patients with pubic arch interference,” Nov 2021. [Online]. Available: https://www.ncbi.nlm.nih.gov/pmc/articles/PMC9838807/

[29] Y. Z. Kaiyu Wu, Bing Li and X. Dai, “Review of research on path planning and control methods of flexible steerable needle puncture robot,” Computer Assisted Surgery, vol. 27, no. 1, pp. 91–112, 2022, pMID: 36052822. [Online]. Available: 10.1080/24699322.2021.2023647

[30] A. Kuntz, M. Emerson, T. E. Ertop, I. Fried, M. Fu, J. Hoelscher, M. Rox, J. Akulian, E. A. Gillaspie, Y. Z. Lee, et al., “Autonomous medical needle steering in vivo,” Science Robotics, vol. 8, no. 82, p. eadf7614, 2023.

[31] K. Seffen, “On the behavior of folded tape-springs,” J. Appl. Mech., vol. 68, no. 3, pp. 369–375, 2001.

[32] K. Seffen and S. Pellegrino, “Deployment of a rigid panel by tape-springs,” Cambridge Univ. Dept. of Engineering, Tech. Rep., July 1997.

[33] K. Seffen and S. Pellegrino, “Deployment dynamics of tape springs,” Proc. of the Royal Society of London. Series A: Mathematical, Physical and Engineering Sciences, vol. 455, no. 1983, pp. 1003–1048, 1999.

[34] W. C. Mitchell, A. Staniforth, and I. Scott, “Analysis of ackermann steering geometry,” SAE Technical Paper, Tech. Rep., 2006.

[35] T. L. De Jong, N. J. Van De Berg, L. Tas, A. Moelker, J. Dankelman, and J. J. Van Den Dobbelsteen, “Needle placement errors: do we need steerable needles in interventional radiology?” Medical Devices: Evidence and Research, pp. 259–265, 2018.

